# Variational Inference of Population Structure in Large SNP Datasets

**DOI:** 10.1101/001073

**Authors:** Anil Raj, Matthew Stephens, Jonathan K. Pritchard

**Affiliations:** Department of Genetics, Stanford University, Stanford, CA, 94305; Departments of Statistics and Human Genetics, University of Chicago, Chicago, IL, 60637; Departments of Genetics and Biology, Howard Hughes Medical Institute, Stanford, CA, 94305

**Keywords:** variational inference, population structure

## Abstract

Tools for estimating population structure from genetic data are now used in a wide variety of applications in population genetics. However, inferring population structure in large modern data sets imposes severe computational challenges. Here, we develop efficient algorithms for approximate inference of the model underlying the STRUCTURE program using a variational Bayesian framework. Variational methods pose the problem of computing relevant posterior distributions as an optimization problem, allowing us to build on recent advances in optimization theory to develop fast inference tools. In addition, we propose useful heuristic scores to identify the number of populations represented in a dataset and a new hierarchical prior to detect weak population structure in the data. We test the variational algorithms on simulated data, and illustrate using genotype data from the CEPH-Human Genome Diversity Panel. The variational algorithms are almost two orders of magnitude faster than STRUCTURE and achieve accuracies comparable to those of ADMIXTURE. Furthermore, our results show that the heuristic scores for choosing model complexity provide a reasonable range of values for the number of populations represented in the data, with minimal bias towards detecting structure when it is very weak. Our algorithm, fastSTRUCTURE, is freely available online at http://pritchardlab.stanford.edu/structure.html.

## INTRODUCTION

Identifying the degree of admixture in individuals and inferring the population of origin of specific loci in these individuals is relevant for a variety of problems in population genetics. Examples include correcting for population stratification in genetic association studies (Pritchard and Donnelly 2001; Price *et al*. 2006), conservation genetics (Wasser *et al*. 2007), and studying the ancestry and migration patterns of natural populations (Rosenberg *et al*. 2002; Reich *et al*.; Catchen *et al*. 2013). With decreasing costs in sequencing and genotyping technologies, there is an increasing need for fast and accurate tools to infer population structure from very large genetic data sets.

Principal components analysis (PCA)-based methods for analyzing population structure, like EIGENSTRAT (Price *et al*. 2006) and SMARTPCA (Patterson *et al*. 2006), construct low-dimensional projections of the data that maximally retain the variance-covariance structure among the sample genotypes. The availability of fast and efficient algorithms for singular value decomposition has enabled PCA-based methods to become the popular choice for analyzing structure in genetic data sets. However, while these low-dimensional projections allow for straightforward visualization of the underlying population structure, it is not straightforward to derive and interpret estimates for global ancestry of sample individuals from their projection coordinates (Novembre and Stephens 2008). In contrast, model-based approaches like STRUCTURE (Pritchard *et al*. 2000) propose an explicit generative model for the data based on the assumptions of Hardy-Weinberg equilibrium between alleles and linkage equilibrium between genotyped loci. Global ancestry estimates are then computed directly from posterior distributions of the model parameters, as done in STRUCTURE, or maximum likelihood estimates of model parameters, as done in FRAPPE (Tang *et al*. 2005) and ADMIXTURE (Alexander *et al*. 2009).

STRUCTURE (Pritchard *et al*. 2000; Falush *et al*. 2003; Hubisz *et al*. 2009) takes a Bayesian approach to estimate global ancestry by sampling from the posterior distribution over global ancestry parameters using a Gibbs sampler that appropriately accounts for the conditional independence relationships between latent variables and model parameters. Simultaneously, the algorithm uses these samples to approximate the log marginal likelihood of the data by a function of the conditional mean and variance of the Bayesian deviance given by the data; this approximation is then used to estimate model complexity (i.e., the number of populations represented in the sample). However, even well-designed sampling schemes need to generate a large number of posterior samples in order to resolve convergence and mixing issues and yield accurate estimates of ancestry proportions, greatly increasing the time complexity of inference for large genotype data sets. To provide faster estimation, FRAPPE and ADMIXTURE both use a maximum likelihood approach. FRAPPE computes maximum likelihood estimates of the parameters of the same model using an expectation-maximization algorithm, while ADMIXTURE computes the same estimates using a sequential quadratic programming algorithm with a quasi-Newton acceleration scheme. Our goal in this paper is to adapt a popular approximate inference framework to greatly speed up inference of population structure while achieving accuracies comparable to STRUCTURE and ADMIXTURE.

Variational Bayesian inference aims to repose the problem of inference as an optimization problem rather than a sampling problem. Variational methods, originally used for approximating intractable integrals, have been used for a wide variety of applications in complex networks (Hofman and Wiggins 2008), machine learning (Jordan *et al*. 1998), (Blei *et al*. 2003) and Bayesian variable selection (Logsdon *et al*. 2010; Carbonetto and Stephens 2012) Variational Bayesian techniques approximate the log marginal likelihood of the data by proposing a family of tractable parametric posterior distributions (variational distribution) over hidden variables in the model; the goal is then to find the optimal member of this family that best approximates the marginal likelihood of the data (see Models and Methods for more details). Thus, a single optimization problem gives us both an approximate estimate of the intractable marginal likelihood and approximate analytical forms for the posterior distributions over unknown variables. Some commonly used optimization algorithms for variational inference include the variational expectation maximization algorithm (Beal 2003), collapsed variational inference (Teh *et al*. 2007), and stochastic gradient descent (Sato 2001).

In the Models and Methods section, we briefly describe the model underlying STRUCTURE and detail the framework for variational Bayesian inference that we use to infer the underlying ancestry proportions. We then propose a more flexible prior distribution over a subset of hidden parameters in the model and demonstrate that estimation of these hyper-parameters using an empirical Bayesian framework improves the accuracy of global ancestry estimates when the underlying population structure is more difficult to resolve. Finally, we describe a scheme to accelerate computation of the optimal variational distributions and describe a set of scores to evaluate the accuracy of the results and to select the number of populations underlying the data. In the Applications section, we compare the accuracy and time complexity of variational inference with those of STRUCTURE and ADMIXTURE on simulated genotype data sets and demonstrate the results of variational inference on a large dataset genotyped in the Human Genome Diversity Panel.

## MODELS AND METHODS

We now briefly describe our generative model for population structure followed by a detailed description of the variational framework used for model inference.

**Variational inference:** Suppose we have *N* diploid individuals genotyped at *L* biallelic loci. A population is represented by a set of allele frequencies at the *L* loci, *P*_*k*_ ∈ [0,1]^*L*^, *k* ∈ {1,…, *K*}, where *K* denotes the number of populations. The allele being represented at each locus can be chosen arbitrarily. Allowing for admixed individuals in the sample, we assume each individual to be represented by a *K*-vector of admixture proportions, *Q*_*n*_ ∈ [0,1]^*K*^, ∑_*k*_ *Q*_*nk*_ = 1, *n* ∈ {1,…, *N*}. Conditioned on *Q*_*n*_, the population assignments of the two copies of a locus, 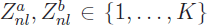, are assumed to be drawn from a multinomial distribution parametrized by *Q*_*n*_. Conditioned on population assignments, the genotype at each locus *G*_*nl*_ is the sum of two independent Bernoulli distributed random variables, each representing the allelic state of each copy of a locus and parameterized by population-specific allele frequencies. The generative process for the sampled genotypes can now be formalized as:

- 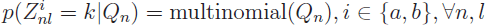
- 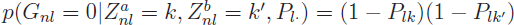
- 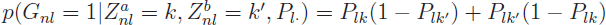
- 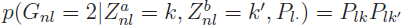

Given the set of sampled genotypes, we can either compute the maximum likelihood estimates of the parameters *P* and *Q* of the model (Alexander *et al*. 2009; Tang *et al*. 2005) or sample from the posterior distributions over the unobserved random variables *Z*^*a*^, *Z*^*b*^, *P*, and *Q* (Pritchard *et al*. 2000) to compute relevant moments of these variables.

Variational Bayesian (VB) inference formulates the problem of computing posterior distributions (and their relevant moments) into an optimization problem. The central aim is to find an element of a tractable family of probability distributions, called variational distributions, that is closest to the true intractable posterior distribution of interest. A natural choice of distance on probability spaces is the Kullback-Leibler (KL) divergence, defined for a pair of probability distributions *q(x)* and *p(x)* as

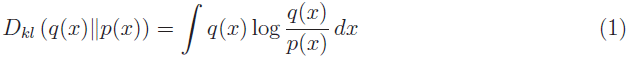

Given the asymmetry of the KL divergence, VB inference chooses *p(x)* to be the intractable posterior and *q(x)* to be the variational distribution; this choice allows us to compute expectations with respect to the tractable variational distribution, often exactly. Except for unrealistically small problem sizes, the KL divergence with respect to the true posterior cannot be computed. However, the KL divergence quantifies the tightness of a lower bound to the log marginal likelihood of the data (Beal 2003); i.e., the KL divergence is equal to the lower bound up to a constant that is a function of only the data and prior parameters. For any variational distribution *q*(*Z*^*a*^, *Z*^*b*^, *P, Q*), we have

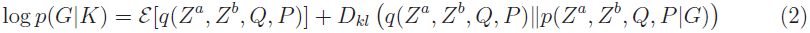

where *ε* is a lower bound to the log marginal likelihood of the data, log *p*(*G*|*K*). An approximation to the true intractable posterior distribution can be computed by minimizing the KL divergence between the true posterior and variational distribution.

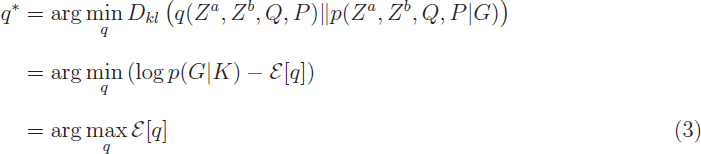

The log marginal likelihood lower bound (LLBO) of the observed genotypes can be written as

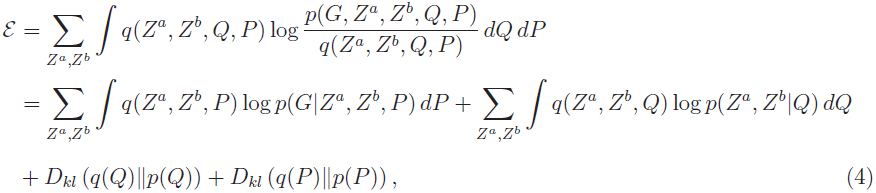

where *p*(*Q*) is the prior on the admixture proportions and *p*(*P*) is the prior on the allele frequencies.

We will restrict our optimization over a variational family that explicitly assumes independence between the latent variables (*Z*^*a*^, *Z*^*b*^) and parameters (*P*, *Q*), commonly called the *mean field approximation* in the statistical physics (Kadanoff 2009) and machine learning literature (Jordan *et al*. 1998)). Since this assumption is certainly not true when inferring population structure, the true posterior will not be a member of the variational family and we will only be able to find the fully factorizable variational distribution that best approximates the true posterior. Nevertheless, this approximation significantly simplifies the optimization problem. Furthermore, we observe empirically that this approximation achieves reasonably accurate estimates of lower order moments (e.g., posterior mean and variance) when the true posterior is replaced by the variational distributions (e.g., when computing prediction error on held-out entries of the genotype matrix). The variational family we choose here is

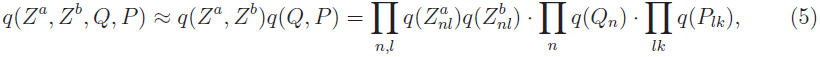

where each factor can then be written as

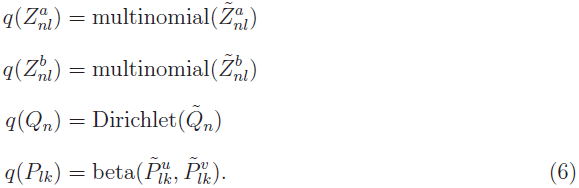

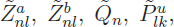 and 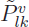 are the parameters of the variational distributions (variational parameters). The choice of the variational family is restricted only by the tractability of computing expectations with respect to the variational distributions; here, we choose parametric distributions that are conjugate to the distributions in the likelihood function. The LLBO of the data in terms of the variational parameters is specified in Appendix-A.

**Priors:** The choice of priors *p*(*Q*_*n*_) and *p*(*P*_*lk*_) plays an important role in inference, particularly when the *F*_*ST*_ between the underlying populations is small and population structure is difficult to resolve. Typical genotype datasets contain hundreds of thousands of genetic variants typed in several hundreds of samples. Given the small sample sizes in these data relative to underlying population structure, the posterior distribution over population allele frequencies can be difficult to estimate; thus, the prior over *P*_*lk*_ plays a more important role in accurate inference than the prior over admixture proportions. Throughout this study, we choose a symmetric Dirichlet prior over admixture proportions; 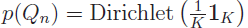.

Depending on the difficulty in resolving structure in a given dataset, we suggest using one of three priors over allele frequencies. When the number of samples is sufficiently large to resolve the underlying population structure, we propose the choice of a flat beta prior over population-specific allele frequencies at each locus; *p*(**P*_*lk*_*) = beta(1,1) (we refer to this prior as “simple prior” throughout the rest of the paper). For genetic data where structure is difficult to resolve, the *F*-model for population structure (Falush *et al*. 2003) proposes a hierarchical prior, based on a demographic model that allows the allele frequencies of the populations to have a shared underlying pattern at all loci. Assuming a star-shaped genealogy where each of the populations simultaneously split from an ancestral population, the allele frequency at a given locus is generated from a beta distribution centered at the ancestral allele frequency at that locus, with variance parametrized by a population-specific drift from the ancestral population (we refer to this prior as “F-prior”).

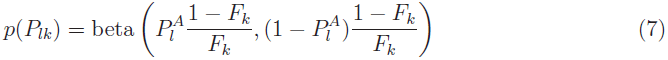

Alternatively, we propose a hierarchical prior that is more flexible than the F-prior and allows for more tractable inference, particularly when additional priors on the hyperparameters need to be imposed. At a given locus, the population-specific allele frequency is generated by a logistic normal distribution, with the normal distribution having a locus-specific mean and a population-specific variance (we refer to this prior as “logistic prior”).

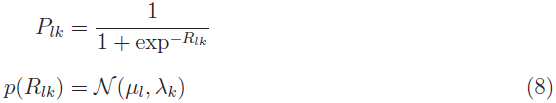

Having specified the appropriate prior distributions, the optimal variational parameters can be computed by iteratively minimizing the KL divergence (or, equivalently, maximizing the LLBO) with respect to each variational parameter, keeping the other variational parameters fixed. The LLBO is concave in each parameter; thus, convergence properties of this iterative optimization algorithm, also called the variational Bayesian expectation maximization algorithm, are similar to those of the expectation-maximization algorithm for maximum likelihood problems. The update equations for each of the three models are detailed in the Appendix-A. Furthermore, when population structure is difficult to resolve, we propose updating the hyperparameters ((*F, P*^*A*^) for the F-prior and (*μ*, λ) for the logistic prior) by maximizing the LLBO with respect to these variables; conditional on these hyperparameter values, improved estimates for the variational parameters are then computed by minimizing the KL divergence. Although such a hyperparameter update is based on optimizing a lower bound on the marginal likelihood, it is likely (although not guaranteed) to increase the marginal likelihood of the data, often leading to better inference. A natural extension of this hierarchical prior would be to allow for a full locus-independent variance-covariance matrix (Pickrell and Pritchard 2012). However, we observed in our simulations that estimating the parameters of this hierarchical prior encouraged the model to strongly overfit the data, leading to poor prediction accuracy on held-out data. Thus, we did not consider this extension in our analyses.

**Accelerated variational inference:** Similar to the EM algorithm, the convergence of the iterative algorithm for variational inference can be quite slow. Treating the iterative update equations for the set of variational parameters 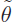 as a deterministic map 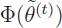, a globally convergent algorithm with improved convergence rates can be derived by adapting the Cauchy-Barzilai-Borwein method for accelerating the convergence of linear fixed-point problems (Raydan and Svaiter 2002) to the nonlinear fixed-point problem given by our deterministic map (Varadhan and Roland 2008). Specifically, given a current estimate of parameters 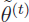, the new estimate can be written as

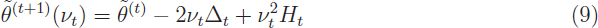

where 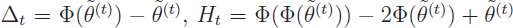 and 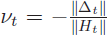. Note that the new estimate is a continuous function of *v*_*t*_ and the standard variational iterative scheme can be obtained from 9 by setting *v*_*t*_ to −1. Thus, for values of *v*_*t*_ close to −1, the accelerated algorithm retains the stability and monotonicity of standard EM algorithms while sacrificing a gain in convergence rate. When *v*_*t*_ < −1, we gain significant improvement in convergence rate, with two potential problems: (a) the LLBO could decrease, i.e., 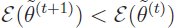, and (b) the new estimate 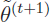 might not satisfy the constraints of the optimization problem. In our experiments, we observe the first problem to occur rarely and we resolve this by simply testing for convergence of the magnitude of difference in LLBO at successive iterations. We resolve the second problem using a simple back-tracking strategy of halving the distance between *v*_*t*_ and −1: *v*_*t*_ ← (*v*_*t*_ −1)/2, until the new estimate 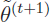 satisfies the constraints of the optimization problem.

**Validation scores:** For each simulated data set, we evaluate the accuracy of each algorithm using two metrics: accuracy of the estimated admixture proportions and the prediction error for a subset of entries in the genotype matrix that are held-out before estimating the parameters. For a given choice of model complexity *K*, an estimate of the admixture proportions *Q\** is taken to be the maximum likelihood estimate of *Q* when using ADMIXTURE, the maximum a posteriori (MAP) estimate of *Q* when using STRUCTURE, and the mean of the variational distribution over *Q* inferred using fastSTRUCTURE. We measure the accuracy of *Q\** by computing the Jensen-Shannon (JS) divergence between *Q\** and the true admixture proportions. The Jensen-Shannon divergence between two probability vectors *P* and *Q* is a bounded distance metric defined as

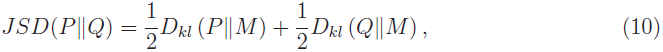

where 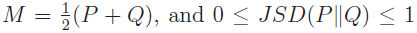. Note that if the lengths of *P* and *Q* are not the same, the smaller vector is extended by appending zero-valued entries. The mean admixture divergence is then defined as the minimum over all permutations of population labels of the mean JS divergence between the true and estimated admixture proportions over all samples, with higher divergence values corresponding to lower accuracy.

We evaluate the prediction accuracy by estimating model parameters (or posterior distributions over them) after holding out a subset *𝒨* of the entries in the genotype matrix. For each held-out entry, the expected genotype is estimated by ADMIXTURE from maximum likelihood parameter estimates as

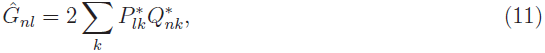

where 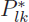 is the maximum likelihood estimate of *P*_*lk*_. The expected genotype given the variational distributions requires integration over the model parameters and is derived in Appendix-B. Given the expected genotypes for the held-out entries, for a specified model complexity *K*, the prediction error is quantified by the deviance residuals under the binomial model averaged over all entries.

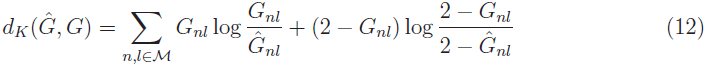

**Model complexity:** ADMIXTURE suggests choosing the value of model complexity *K* that achieves the smallest value of *d*_*K*_(*Ĝ, G*), i.e., 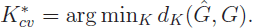. We propose two additional metrics to select model complexity in the context of variational Bayesian inference. Assuming a uniform prior on *K*, the optimal model complexity 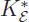 is chosen to be the one that maximizes the LLBO, where the LLBO is used as an approximation to the marginal likelihood of the data. However, since the difference between the log marginal likelihood of the data and the LLBO is difficult to quantify, the trend of LLBO as a function of *K* cannot be guaranteed to match that of the log marginal likelihood. Additionally, we propose a useful heuristic to choose *K* based on the tendency of mean-field variational schemes to populate only those model components that are essential to explain patterns underlying the observed data. Specifically, given an estimate of *Q\** obtained from variational inference executed for a choice of *K*, we compute the ancestry contribution of each model component as the mean admixture proportion over all samples, i.e., 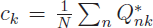. The number of relevant model components 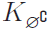 is then the minimum number of populations that have a cumulative ancestry contribution of at least 99.99%.

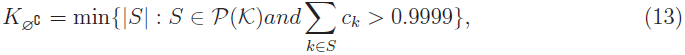

where *𝒦* = {1, …, *K*} and 𝒫(𝒦) is the power set of 𝒦. As *K* increases, 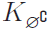 tends to approach a limit that can be chosen as the optimal model complexity 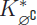.

## APPLICATIONS

In this section, we compare the accuracy and runtime performance of the variational inference framework with the results of STRUCTURE and ADMIXTURE both on datasets generated from the *F*-model and on the Human Genome Diversity Panel (HGDP) (Rosenberg *et al*. 2002). We expect the results of ADMIXTURE to match those of FRAPPE (Tang *et al*. 2005) since they both compute maximum likelihood estimates of the model parameters. However, ADMIXTURE converges faster than FRAPPE allowing us to compare it with fastSTRUCTURE using thousands of simulations. In general, we observe that fastSTRUCTURE estimates ancestry proportions with accuracies comparable to, and sometimes better than, those estimated by ADMIXTURE even when the underlying population structure is rather weak. Furthermore, fastSTRUCTURE is about two orders of magnitude faster than STRUCTURE and has comparable runtimes to that of ADMIXTURE. Finally, fastSTRUCTURE gives us a reasonable range of values for the model complexity required to explain structure underlying the data, without the need for a cross-validation scheme. Below, we highlight the key advantages and disadvantages of variational inference in each problem setting.

**Simulated datasets:** To evaluate the performance of the different learning algorithms, we generated two groups of simulated genotype datasets, with each genotype matrix consisting of 600 samples and 2500 loci. The first group was used to evaluate the accuracy of the algorithms as a function of strength of the underlying population structure while the second group was used to evaluate accuracy as a function of number of underlying populations. Although the size of each genotype matrix was kept fixed in these simulations, the performance characteristics of the algorithms are expected to be similar if the strength of population structure is kept fixed and the dataset size is varied (Patterson *et al*. 2006).

For the first group, the samples were drawn from a 3-population demographic model as shown in Figure 1a. The edge weights correspond to the parameter *F* in the model that quantifies the genetic drift of each of the three current populations from an ancestral population. We introduced a scaling factor *r* ∈ [0,1] that quantifies the resolvability of population structure underlying the samples. Scaling *F* by *r* reduces the amount of drift of current populations from the ancestral population; thus, structure is difficult to resolve when *r* is close to 0, while structure is easy to resolve when *r* is close to 1. For each *r* ∈ {0.05, 0.10, …, 0.95, 1}, we generated 50 replicate datasets. The ancestral allele frequencies *π*^*A*^ for each dataset were drawn from the frequency spectrum computed using the HGDP panel to simulate allele frequencies in natural populations. For each dataset, the allele frequency at a given locus for each population was drawn from a beta distribution with mean 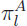 and variance 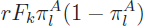, and the admixture proportions for each sample were drawn from a symmetric Dirichlet distribution, namely Dirichlet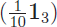, in order to simulate small amounts of gene flow between the three populations. Finally, 10% of the samples in each dataset, randomly selected, were assigned to one of the three populations with zero admixture.

For the second group, the samples were drawn from a star-shaped demographic model with *K*_*t*_ populations. Each population was assumed to have equal drift from an ancestral population, with the F parameter fixed at either 0.01 to simulate weak structure or 0.04 to simulate strong structure. The ancestral allele frequencies were simulated similar to the first group and 50 replicate datasets were generated for this group for each value of *K*_*t*_ ∈ {1,…, 5}. We executed ADMIXTURE and fastSTRUCTURE for each dataset with various choices of model complexity; for datasets in the first group, model complexity *K* ∈ {1, …, 5}, and for those in the second group *K* ∈{1, …, 8}. We executed ADMIXTURE with default parameter settings; with these settings the algorithm terminates when the increase in log likelihood is less than 10^−4^ and computes prediction error using 5-fold cross-validation. fastSTRUCTURE was executed with a convergence criterion of change in the per-genotype log marginal likelihood lower bound |Δ *ε*| < 10^−8^. We held out 20 random disjoint genotype sets each containing 1% of entries in the genotype matrix and used the mean and standard error of the deviance residuals for these held-out entries as an estimate of the prediction error.

For each group of simulated datasets, we illustrate a comparison of the performance of ADMIXTURE and fastSTRUCTURE with the simple and the logistic prior. When structure was easy to resolve, both the F-prior and the logistic prior returned similar results; however, the logistic prior returned more accurate ancestry estimates when structure was difficult to resolve. Plots including results using the F-prior are shown in the supplementary figures. Since ADMIXTURE uses held-out deviance residuals to choose model complexity, we demonstrate the results of the two algorithms, each using deviance residuals to choose *K*, using solid lines in Figures 1 and 2. Additionally, in these figures, we also illustrate the performance of fastSTRUCTURE, when using the two alternative metrics to choose model complexity, using blue lines.

**Figure 1.**
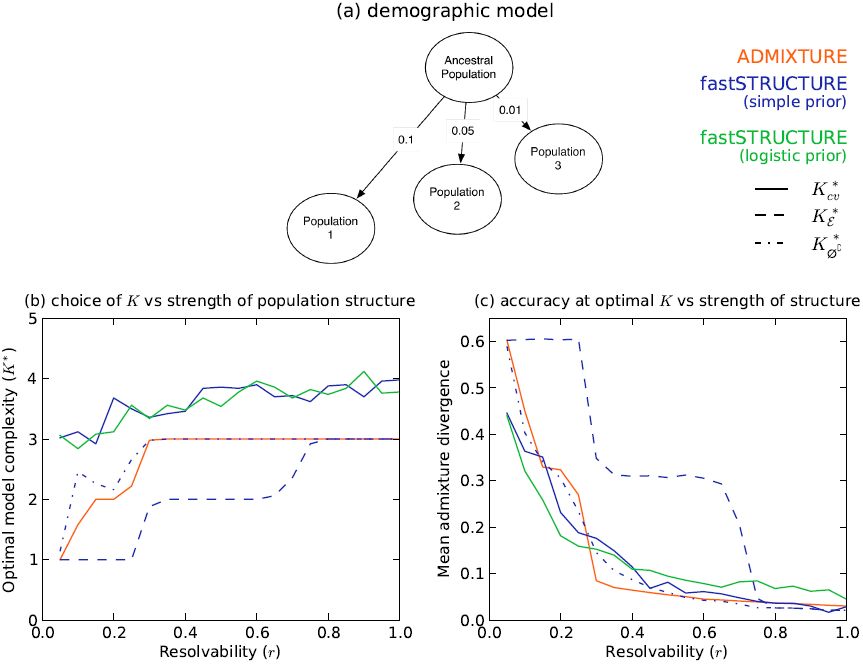
Accuracy of different algorithms as a function of resolvability of population structure. Subfigure (a) illustrates the demographic model underlying the three populations represented in the simulated datasets. The edge weights quantify the amount of drift from the ancestral population. In (b) and (c), ‘Resolvability’ is a scalar by which the population-specific drifts in the demographic model are multiplied, with higher values of resolvability corresponding to stronger structure. Subfigure (b) compares the optimal model complexity given the data, averaged over 50 replicates, inferred by ADMIXTURE 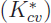, fastSTRUCTURE with simple prior 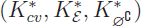, and fastSTRUCTURE with logistic prior 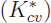. Subfigure (c) compares the accuracy of admixture proportions, averaged over replicates, estimated by each algorithm at the optimal value of *K* in each replicate.

**Figure 2.**
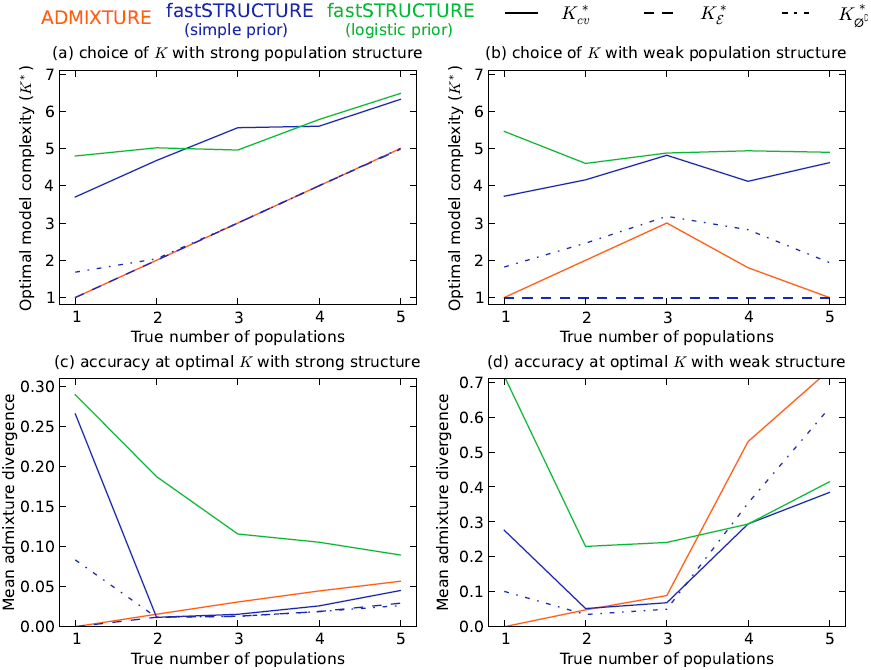
Accuracy of different algorithms as a function of the true number of populations. The demographic model is a star-shaped genealogy with populations having undergone equal amounts of drift. The left and right panels correspond to strong structure (*F* = 0.04) and weak structure (*F* = 0.01), respectively. Subfigures in the top panel compare the optimal model complexity estimated by the different algorithms using various metrics, averaged over 50 replicates, to the true number of populations represented in the data. Notably, when population structure is weak, both ADMIXTURE and fastSTRUCTURE fail to detect structure when the number of populations is too large. Subfigures in the bottom panel compare the accuracy of admixture proportions estimated by each algorithm at the optimal model complexity for each replicate.

**Choice of** *K*: Identifying the number of populations needed to explain structure in the data is a problem of great interest associated with the inference of population structure. While this commonly has been addressed using cross-validation or a model-selection framework, the problem of identifying a single “correct” number of populations is ill-posed and strongly dependent on how pertinent the underlying model of population structure is to a specific study sample (Engelhardt and Stephens 2010). For example, given a set of individuals sampled from a habitat with spatially continuous population structure, applying ADMIXTURE or STRUCTURE to the sample genotypes would give us insights into population structure represented in the data. However, the number of populations returned by an automatic scheme to select *K* is not likely to be meaningful in this case, and could be strongly dependent on the ascertainment of individuals in the dataset and the deviation of sample genotypes from the strict random-mating model. While identifying a reasonable range of values for *K* for a given dataset is certainly useful, the specific values of *K* and the identified populations need to be interpreted within the context of prior knowledge specific to the dataset being analyzed.

The manual of the ADMIXTURE code proposes choosing model complexity that minimizes the prediction error on held-out data estimated using the mean deviance residuals reported by the algorithm 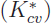. In Figure 1b, using the first group of simulations, we compare the value of 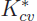, averaged over 50 replicate datasets, between the two algorithms as a function of the resolvability of population structure in the data. We observe that while deviance residuals estimated by ADMIXTURE robustly identify an appropriate model complexity, the value of *K* identified using deviance residuals computed using the variational parameters from fastSTRUCTURE appear to over-estimate the value of *K* underlying the data. However, on closer inspection, we observe that the difference in prediction errors between large values of *K* are statistically insignificant (Figure 3, middle panels). This suggests the following heuristic: select the lowest model complexity above which prediction errors don’t vary significantly.

**Figure 3.**
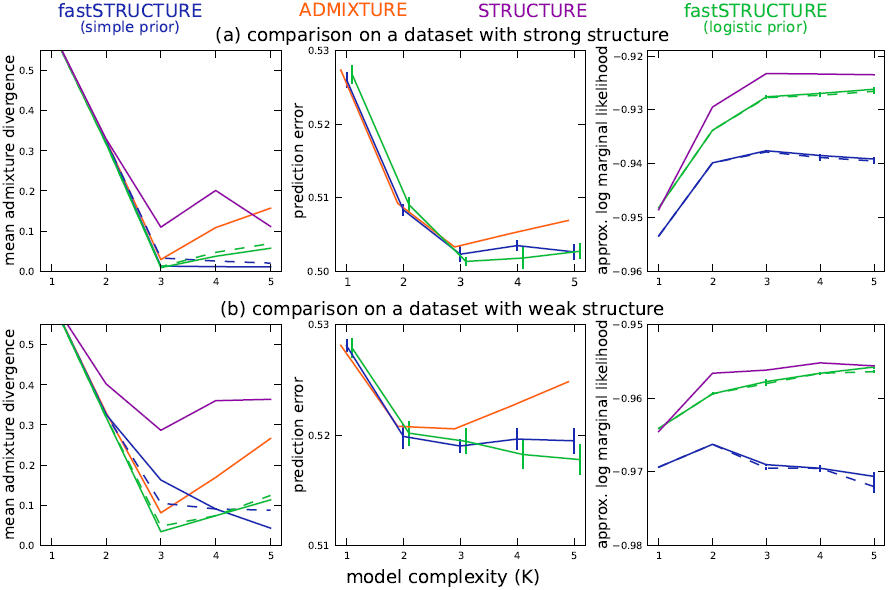
Accuracy of different algorithms as a function of model complexity (*K*) on two simulated data sets, one in which ancestry is easy to resolve (top panel; *r* = 1) and one in which ancestry is difficult to resolve (bottom panel; *r* = 0.5). Solid lines correspond to parameter estimates computed with a convergence criterion of |Δ*ε*| < 10^−8^, while the dashed lines correspond to a weaker criterion of |Δ*ε*| < 10^−6^. The left panel of subfigures shows the mean admixture divergence between the true and inferred admixture proportions while the middle panel shows the mean binomial deviance of held-out genotype entries. Note that for values of *K* greater than the optimal value, any change in prediction error lies within the standard error of estimates of prediction error suggesting that we should choose the smallest value of model complexity above which a decrease in prediction error is statistically insignificant. The right panel shows the approximations to the marginal likelihood of the data computed by STRUCTURE and fastSTRUCTURE.

Alternatively, for fastSTRUCTURE with the simple prior, we propose two additional metrics for choosing model complexity: (1) 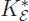: value of *K* that maximizes the LLBO of the entire dataset, and (2) 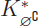: the limiting value, as *K* increases, of the smallest number of model components that accounts for almost all of the ancestry in the sample. In Figure 1b, we observe that 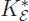 has the attractive property of robustly identifying strong structure underlying the data, while 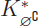 identifies additional model components needed to explain weak structure in the data, with a slight upward bias in complexity when the underlying structure is extremely difficult to resolve. For the second group of simulations, similar to results observed for the first group, when population structure is easy to resolve, ADMIXTURE robustly identifies the correct value of *K* (shown in Figure 2a). However, for similar reasons as before, the use of prediction error with fastSTRUCTURE tends to systematically overestimate the number of populations underlying the data. In contrast, 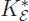 and 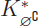 match exactly to the true *K* when population structure is strong. When the underlying population structure is very weak, 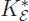 is a severe underestimate of the true *K* while 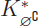 slightly overestimates the value of *K*. Surprisingly, 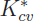 estimated using ADMIXTURE and 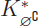 estimated using fastSTRUCTURE tend to underestimate the number of populations when the true number of populations *K*_*t*_ is large, as shown in Figure 2b.

For a new dataset, we suggest executing fastSTRUCTURE for multiple values of *K* and estimating 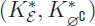 to obtain a reasonable range of values for the number of populations that would explain structure in the data, under the given model. To look for subtle structure in the data, we suggest executing fastSTRUCTURE with the logistic prior with values for values of *K* similar to those identified by using the simple prior.

**Accuracy of ancestry proportions:** We evaluated the accuracy of the algorithms by comparing the divergence between the true admixture proportions and the estimated admixture proportions at the optimal model complexity computed using the above metrics for each dataset. In Figure 1c, we plot the mean divergence between the true and estimated admixture proportions, over multiple replicates, as a function of resolvability. We observe that the admixture proportions estimated by fastSTRUCTURE at 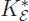 have high divergence; however, this is a result of LLBO being too conservative in identifying *K*. At 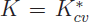 and 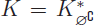, fastSTRUCTURE estimates admixture proportions with accuracies comparable to, and sometimes better than, ADMIXTURE even when the underlying population structure is rather weak. Furthermore, the held-out prediction deviances computed using posterior estimates from variational algorithms are consistently smaller than those estimated by ADMIXTURE (see Figure S3) demonstrating the improved accuracy of variational Bayesian inference schemes over maximum likelihood methods. Similarly, for the second group of simulated datasets, we observe in Figures 2c and 2d that the accuracy of variational algorithms tend to be comparable to or better than that of ADMIXTURE, particularly when structure is difficult to resolve. When structure is easy to resolve, the increased divergence estimates of fastSTRUCTURE with the logistic prior result from the upward bias in the estimate of 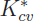; this can be improved by using cross-validation more carefully in choosing model complexity.

**Visualizing ancestry estimates:** Having demonstrated the performance of fastSTRUCTURE on multiple simulated datasets, we now illustrate the performance characteristics and parameter estimates using two specific datasets (selected from the first group of simulated datasets), one with strong population structure (*r* = 1) and one with weak structure (*r* = 0.5). In addition to these algorithms, we executed STRUCTURE for these two datasets using the model of independent allele frequencies in order to directly compare with the results of fastSTRUCTURE. For each dataset, *α* was kept fixed to 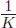 for all populations, similar to the prior used for fastSTRUCTURE and each run consisted of 50000 burn-in steps and 50000 MCMC steps. In Figure 3, we illustrate the divergence of admixture estimates and the prediction error on held-out data each as a function of *K*. For all choices of *K* greater than or equal to the true value, the accuracy of the fastSTRUCTURE, measured using both admixture divergence and prediction error, is generally comparable to or better than that of ADMIXTURE and STRUCTURE, even when the underlying population structure is rather weak. In the right panels of Figure 3, we plot the approximate marginal likelihood of the data, reported by STRUCTURE, and the optimal LLBO, computed by fastSTRUCTURE, each as a function of *K*. We note that the looseness of the bound between STRUCTURE and fastSTRUCTURE can make the LLBO a less reliable measure to choose model complexity than the approximate marginal likelihood reported by STRUCTURE, particularly when the size of the dataset is not sufficient to resolve the underlying population structure.

Figure 4 illustrates the admixture proportions estimated by the different algorithms on both data sets at two values of *K*, using Distruct plots (Rosenberg 2004). For the larger choice of model complexity, we observe that fastSTRUCTURE with the simple prior uses only those model components that are necessary to explain the data, allowing for automatic inference of model complexity (Mackay 2003). To better illustrate this property of unsupervised Bayesian inference methods, the right panels of Figure 4 show the mean contribution of ancestry from each model component to samples in the dataset. While ADMIXTURE uses all components of the model to fit the data, STRUCTURE and fastSTRUCTURE assign negligible posterior mass to model components that are not required to capture structure in the data. The number of non-empty model components (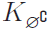) automatically identifies the model complexity required to explain the data; the optimal model complexity 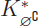 is then the mode of all values of 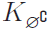 computed for different choices of *K*.

**Figure 4.**
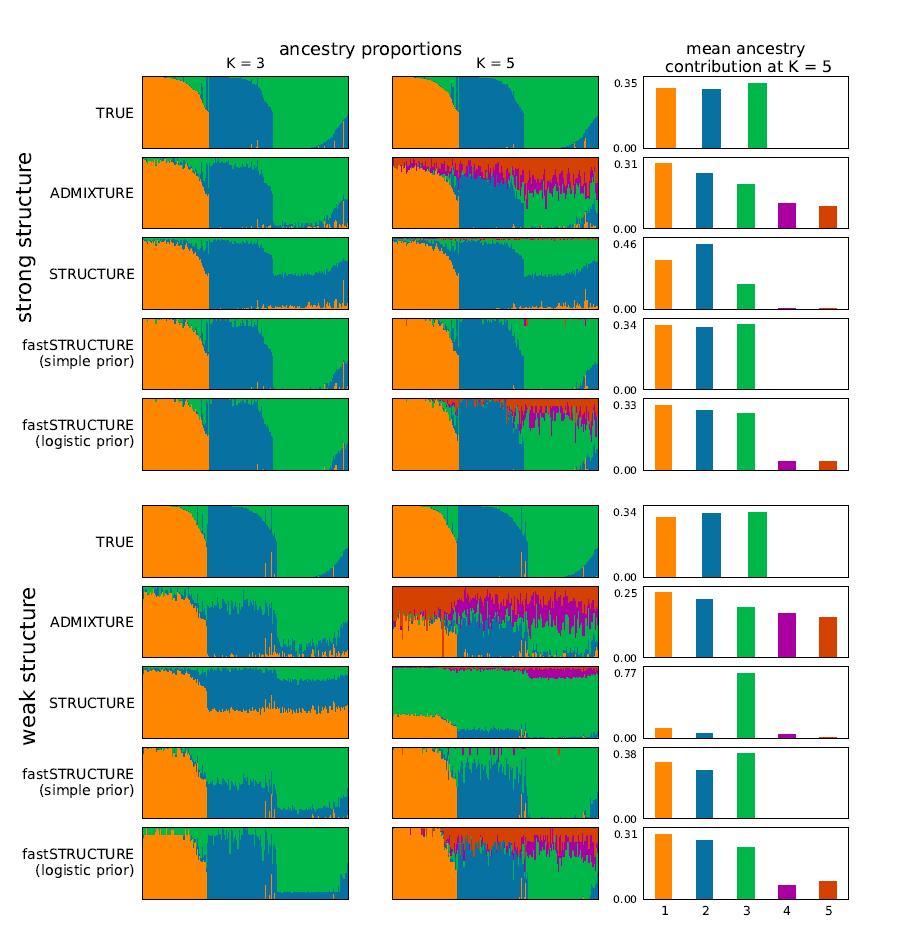
Visualizing ancestry proportions estimated by different algorithms on two simulated data sets, one with strong structure (top panel; *r* = 1) and one with weak structure (bottom panel; *r* = 0.5). For the left and middle panels of subfigures, ancestry estimated at model complexity of *K* = 3 and *K* = 5, respectively, are illustrated. The subpanels within a panel illustrate the true ancestry and the ancestry inferred by each algorithm. Each color represents a population and each individual is represented by a vertical line partitioned into colored segments whose lengths represent the admixture proportions from *K* populations. In the right panel of subfigures, the mean ancestry contributions of the model components are shown, when the model complexity *K* = 5.

When population structure is difficult to resolve, imposing a logistic prior and estimating its parameters using the data is likely to increase the power to detect weak structure. However, estimation of the hierarchical prior parameters by maximizing the approximate marginal likelihood also makes the model susceptible to overfitting by encouraging a small set of samples to be randomly, and often confidently, assigned to unnecessary components of the model. To correct for this, when using the logistic prior, we suggest estimating the variational parameters with multiple random restarts and using the mean of the parameters corresponding to the top 5 values of LLBO. In order to ensure consistent population labels when computing the mean, we permuted the labels for each set of variational parameter estimates to find the permutation with the lowest pairwise Jensen-Shannon divergence between admixture proportions among pairs of restarts. Admixture estimates computed using this scheme show improved robustness against overfitting, as illustrated in Figure 4. Moreover, the pairwise Jensen-Shannon divergence between admixture proportions among all restarts of the variational algorithms can also be used as a measure of the robustness of their results and as a signature of how strongly they overfit the data.

**Runtime performance:** A key advantage of variational Bayesian inference algorithms compared to inference algorithms based on sampling is the dramatic improvement in time complexity of the algorithm. To evaluate the runtimes of the different learning algorithms, we generated from the *F*-model datasets with sample sizes *N* ∈ {200, 600} and numbers of loci *L* ∈ {500, 2500}, each having 3 populations with *r* = 1. The time complexity of each of the above algorithms is linear in the number of samples, loci, and populations, i.e. O(*NLK*); in comparison, the time complexity of principal components analysis is quadratic in the number of samples and linear in the number of loci. In Figure 5, the mean runtime of the different algorithms is shown as a function of problem size defined as *N* × *L* × *K*. The added complexity of the cost function being optimized in fastSTRUCTURE increases its runtime when compared to ADMIXTURE. However, fastSTRUCTURE is about two orders of magnitude faster than STRUCTURE, making it suitable for large datasets with hundreds of thousands of genetic variants. For example, using a dataset with 1000 samples genotyped at 500, 000 loci with *K* = 10, each iteration of our current Python implementation of fastSTRUCTURE with the simple prior takes about 11 minutes, while each iteration of ADMIXTURE takes about 16 minutes. Since one would usually like to estimate the variational parameters for multiple values of *K* for a new dataset, a faster algorithm that gives an approximate estimate of ancestry proportions in the sample would be of much utility, particularly to guide an appropriate choice of *K*. We observe in our simulations that a weaker convergence criterion of |Δ*ε*| < 10^−6^ gives us comparably accurate results with much shorter run times, illustrated by the dashed lines in Figures 3 and 5. Based on these observations, we suggest executing multiple random restarts of the algorithm with a weak convergence criterion of |Δ*ε*| < 10^−5^ to rapidly obtain reasonably accurate estimates of the variational parameters, prediction errors and ancestry contributions from relevant model components.

**Figure 5.**
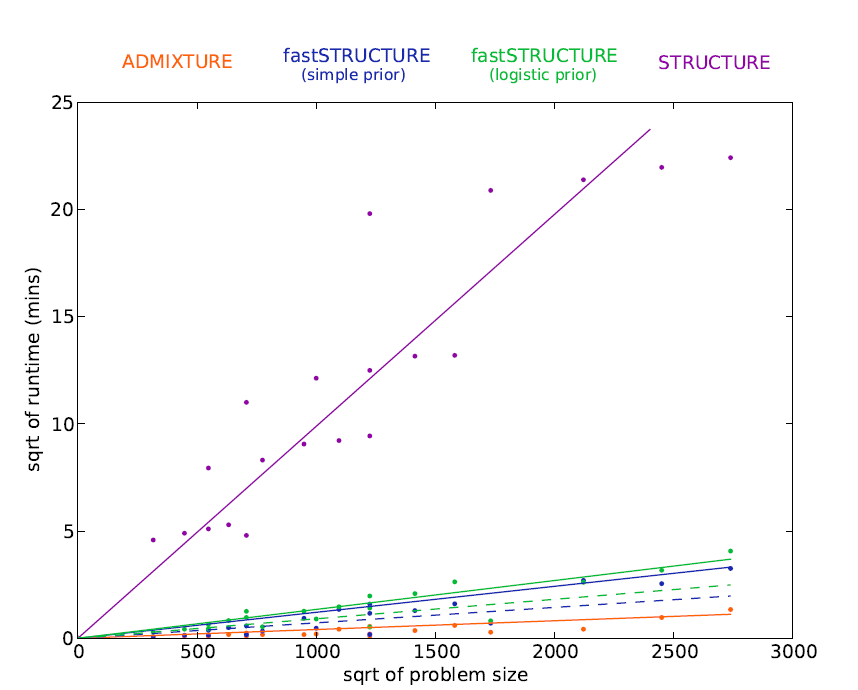
Runtimes of different algorithms on simulated data sets with different number of loci and samples; the square root of runtime (in minutes) is plotted as a function of square root of problem size (defined as *N × *L* × K*). Similar to figure 3, dashed lines correspond to a weaker convergence criterion than solid lines.

**HGDP Panel:** We now compare the results of ADMIXTURE and fastSTRUCTURE on a large, well-studied dataset of genotypes at single nucleotide polymorphisms (SNP) genotyped in the Human Genome Diversity Panel (HGDP) (Li *et al*. 2008), in which 1048 individuals from 51 different populations were genotyped using Illumina’s HumanHap650Y platform. We used the set of 938 “unrelated” individuals for the analysis in this paper. For the selected set of individuals, we removed SNPs that were monomorphic, had missing genotypes in more than 5% of the samples and failed the Hardy-Weinberg Equilibrium (HWE) test at *p* < 0.05 cutoff. To test for violations from HWE, we selected three population groups that have relatively little population structure (East Asia, Europe, Bantu Africa), constructed three large groups of individuals from these populations, and performed a test for HWE for each SNP within each large group. The final dataset contained 938 samples with genotypes at 657,143 loci, with 0.1% of the entries in the genotype matrix missing. We executed ADMIXTURE and fastSTRUCTURE using this dataset with allowed model complexity *K* ∈ {5, …, 15}. In Figure 6, the ancestry proportions estimated by ADMIXTURE and fastSTRUCTURE at *K* = 7 are shown; this value of *K* was chosen to compare with results reported using the same dataset with FRAPPE (Li *et al*. 2008). In contrast to results reported using FRAPPE, we observe that both ADMIXTURE and fastSTRUCTURE identify the Mozabite, Bedouin, Palestinian, and Druze populations as very closely related to European populations with some gene flow from Central-Asian and African populations; this result was robust over multiple random restarts of each algorithm. Since both ADMIXTURE and FRAPPE maximize the same likelihood function, the slight difference in results is likely due to differences in the modes of the likelihood surface to which the two algorithms converge. A notable difference between ADMIXTURE and fastSTRUCTURE is in their choice of the 7^th^ population – ADMIXTURE splits the Native American populations along a north-south divide while fastSTRUCTURE splits the African populations into central African and south African population groups.

**Figure 6.**
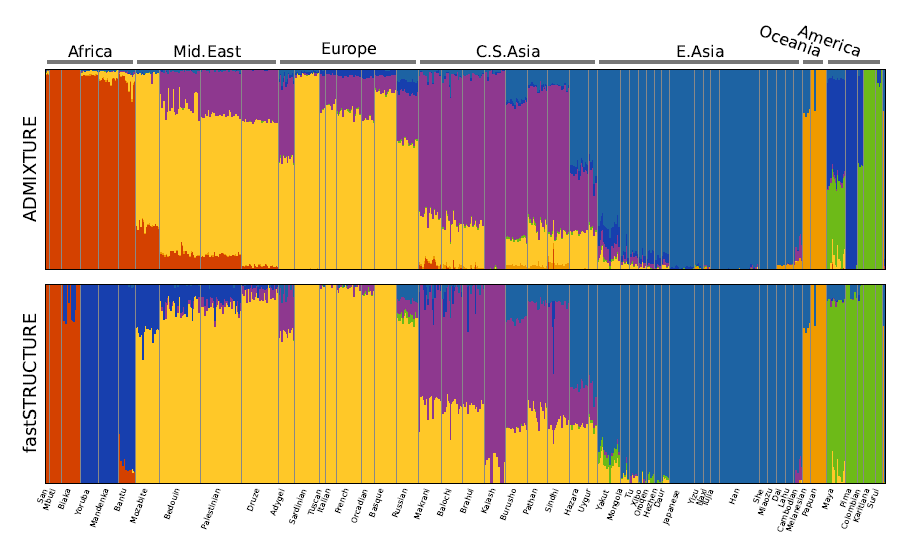
Ancestry proportions inferred by ADMIXTURE and fastSTRUCTURE (with the simple prior) on the HGDP data at *K* = 7 (Li *et al.* 2008). Notably, ADMIXTURE splits the Central and South American populations into two groups while fastSTRUCTURE assigns higher approximate marginal likelihood to a split of sub-Saharan African populations into two groups.

Interestingly, both algorithms strongly suggest the existence of additional weak population structure underlying the data, as shown in Figure 7. ADMIXTURE, using cross-validation, identifies the optimal model complexity to be 11; however, the deviance residuals appear to change very little beyond *K* = 7 suggesting that the model components identified at *K* = 7 explain most of the structure underlying the data. Using LLBO, fastSTRUCTURE identifies the number of easily resolvable populations to be 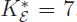, while the 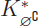 estimate suggests the number of populations to be 9. The lowest cross-validation error for fastSTRUCTURE is achieved at 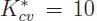; a slight overestimate compared to the model complexity range suggested by 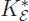 and 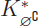.

**Figure 7.**
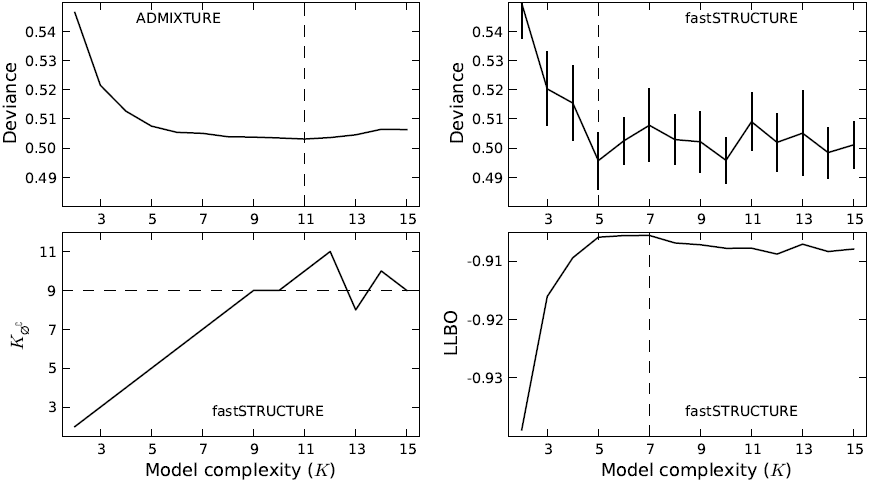
Model choice of ADMIXTURE and fastSTRUCTURE (with the simple prior) on the HGDP data. Optimal value of *K*, identified by ADMIXTURE using deviance residuals, and by fastSTRUCTURE using deviance, 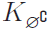, and LLBO, are shown by a dashed line.

The admixture proportions estimated at the optimal choices of model complexity using the different metrics are shown in Figure 8. The admixture proportions estimated at *K* = 7 and *K* = 9 are remarkably similar with the Kalash and Karitiana populations being assigned to their own model components at *K* = 9. These results demonstrate the ability of LLBO to identify strong structure underlying the data and that of 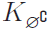 to identify additional weak structure that explain variation in the data. At *K* = 10 (as identified using cross-validation), we observe that only 9 of the model components are populated. However, the estimated admixture proportions differ crucially with all African populations grouped together, the Melanesian and Papuan populations each assigned to their own groups, and the Middle-Eastern populations represented as predominantly an admixture of Europeans and a Bedouin sub-population with small amounts of gene flow from Central-Asian populations.

**Figure 8.**
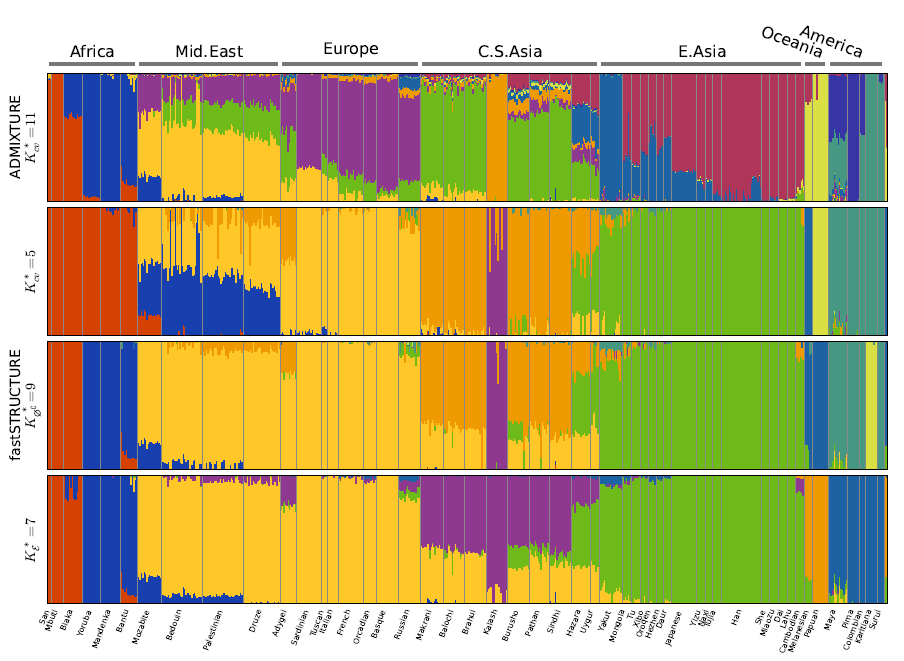
Ancestry proportions inferred by ADMIXTURE and fastSTRUCTURE (with the simple prior) at the optimal choice of *K* identified by relevant metrics for each algorithm. Notably, the admixture proportions at 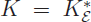 and 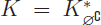 are quite similar, with estimates in the latter case identifying the Kalash and Karitiana as additional separate groups that share very little ancestry with the remaining populations.

## DISCUSSION

Our analyses on simulated and natural datasets demonstrate that fastSTRUCTURE estimates approximate posterior distributions on ancestry proportions two orders of magnitude faster than STRUCTURE, with ancestry estimates and prediction accuracies that are comparable to those of ADMIXTURE. Posing the problem of inference in terms of an optimization problem allows us to draw on powerful tools in convex optimization and plays an important role in the gain in speed achieved by variational inference schemes, when compared to the Gibbs sampling scheme used in STRUCTURE. In addition, the flexible logistic prior enables us to resolve subtle structure underlying a dataset. The considerable improvement in runtime with comparable accuracies allows the application of these methods to large genotype datasets that are steadily becoming the norm in studies of population history, genetic association with disease, and conservation biology.

The choice of model complexity, or the number of populations required to explain structure in a dataset, is a difficult problem associated with the inference of population structure. Unlike in maximum likelihood estimation, the model parameters have been integrated out in variational inference schemes and optimizing the KL divergence in fastSTRUCTURE does not run the risk of overfitting. The heuristic scores that we have proposed to identify model complexity provide a robust and reasonable range for the number of populations underlying the dataset, without the need for a time-consuming cross-validation scheme.

As in the original version of STRUCTURE, the model underlying fastSTRUCTURE does not explicitly account for linkage disequilibrium (LD) between genetic markers. While LD between genotype markers in the genotype dataset will lead us to underestimate the variance of the approximate posterior distributions, the improved accuracy in predicting held-out genotypes for the HGDP dataset demonstrates that the underestimate due to un-modeled LD and the mean field approximation is not too severe. Furthermore, not accounting for LD appropriately can lead to significant biases in local ancestry estimation, depending on the sample size and population haplotype frequencies. However, we believe global ancestry estimates are likely to incur very little bias due to un-modeled LD.

In summary, we have presented a variational framework for fast, accurate inference of global ancestry of samples genotyped at a large number of genetic markers. For a new dataset, we recommend executing our program, fastSTRUCTURE, for multiple values of *K* to obtain a reasonable range of values for the appropriate model complexity required to explain structure in the data, as well as ancestry estimates at those model complexities. For improved ancestry estimates and to identify subtle structure, we recommend executing fastSTRUCTURE with the logistic prior at values of *K* similar to those identified when using the simple prior. Our program is available for download at http://pritchardlab.stanford.edu/structure.html.

## APPENDIX-A

Given the parametric forms for the variational distributions and a choice of prior for the fastSTRUCTURE model, the per-genotype LLBO is given as

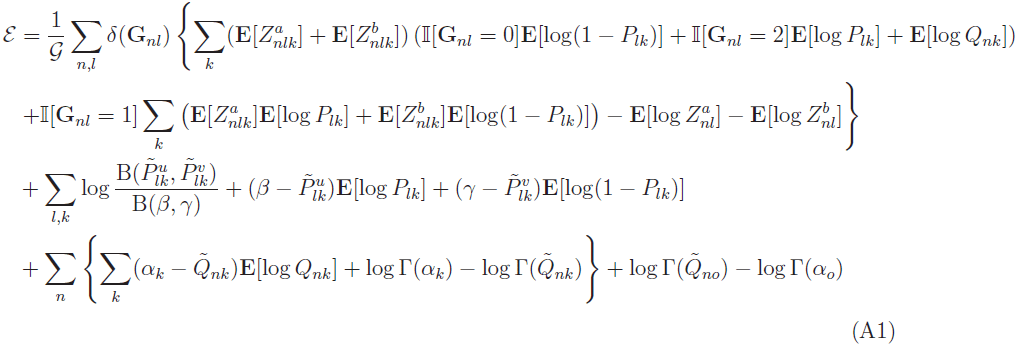

where **E[·]** is the expectation taken with respect to the appropriate variational distribution, B(·) is the beta function, Γ(·) is the gamma function, {*α, β, γ*} are the hyperparameters in the model, *δ*(·) is an indicator variable that takes the value of zero if the genotype is missing, *𝒢* is the number of observed entries in the genotype matrix, *α*_*o*_ = ∑_*k*_*α*_*k*_, and 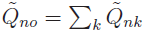. Maximizing this lower bound for each variational parameter, keeping the other parameters fixed, gives us the following update equations.

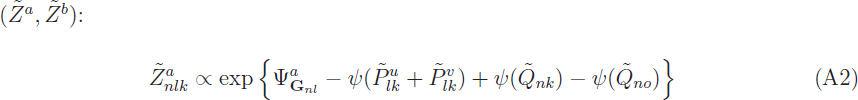

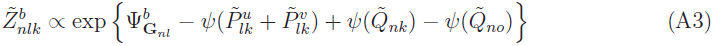

where

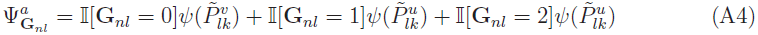

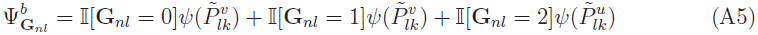

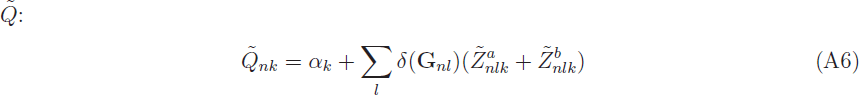

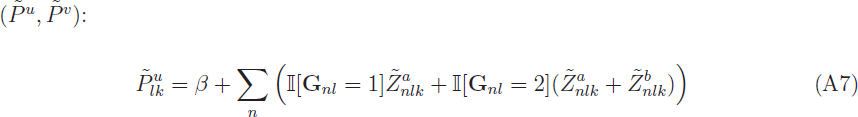

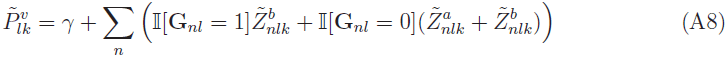

In the above update equations, *ψ*(·) is the digamma function. When the F-prior is used, the LLBO and the update equations remain exactly the same, after replacing *β* with 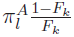 and *γ* with 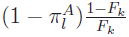. In this case, the LLBO is also maximized with respect to the hyperparameter *F* using the L-BFGS-B algorithm, a quasi-Newton code for bound-constrained optimization.

When the logistic prior is used, a straightforward maximization of the LLBO no longer gives us explicit update equations for 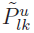 and 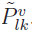. One alternative is to use a constrained optimization solver, like L-BFGS-B; however, the large number of variational parameters to be optimized greatly increases the per-iteration computational cost of the inference algorithm. Instead, we propose update equations for 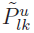 and 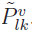 to have a similar form as those obtained with the simple prior,

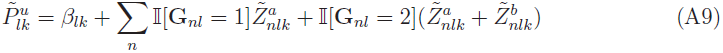

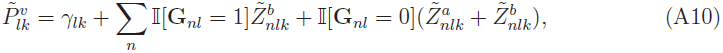

where *β*_*lk*_ and *γ*_*lk*_ implicitly depend on 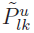 and 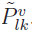 as follows:

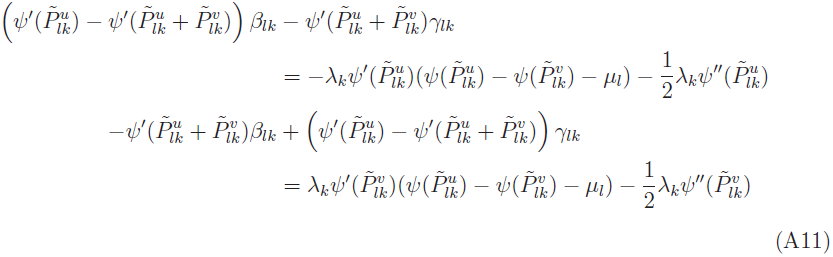

The optimal values for 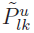 and 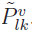 can be obtained by iterating between the two sets of equations to convergence. Thus, when the logistic prior is used, the algorithm is implemented as a nested iterative scheme where for each update of all the variational parameters, there is an iterative scheme that computes the update for 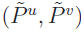. Finally, the optimal value of the hyperparameter *μ* is obtained straightforwardly as

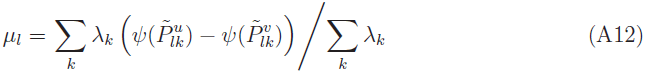

while the optimal λ is computed using a constrained optimization solver.

APPENDIX-B

Given the observed genotypes **G,** the probability of the unobserved genotype for the *n*^*th*^ sample at the *l*^th^ locus is given as

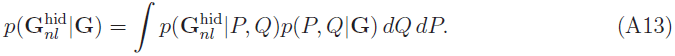

Replacing the posterior *p*(*P, Q***|G)** with the optimal variational posterior distribution, we get

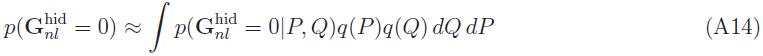

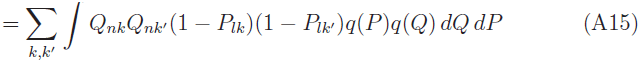

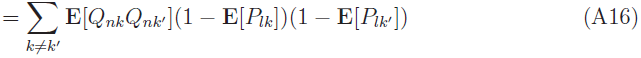

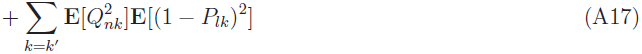

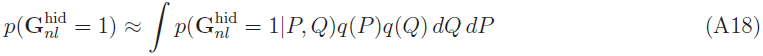

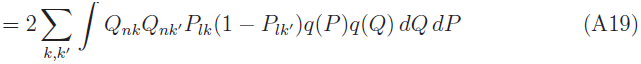

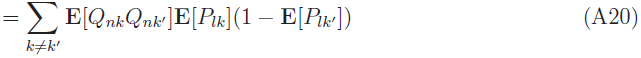

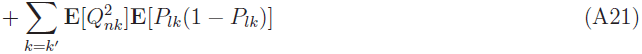

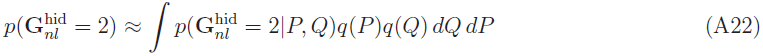

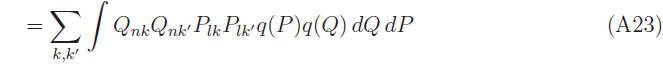

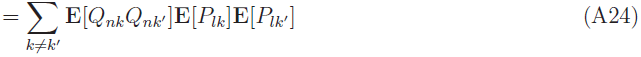

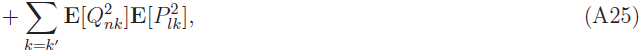

where

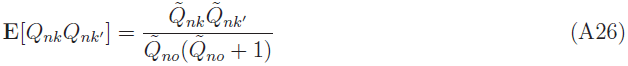

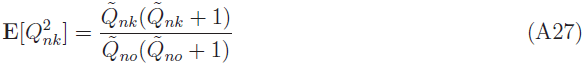

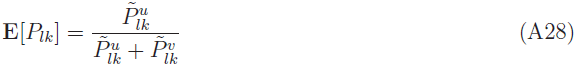

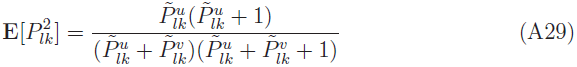

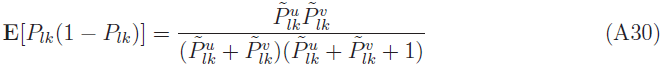

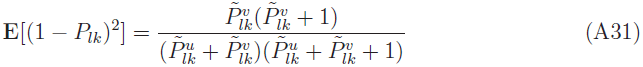

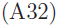

The expected genotype can then be straightforwardly computed from these genotype probabilities.

## LITERATURE CITED

Alexander, D. H., J. Novembre, and K. Lange, 2009 Fast model-based estimation of ancestry in unrelated individuals. Genome Research 19(9): 1655–1664.

Beal, M. J., 2003 Variational Algorithms for Approximate Bayesian Inference. Ph. D. thesis, Gatsby Computational Neuroscience Unit, University College London.

Blei, D. M., A. Y. Ng, and M. I. Jordan, 2003 Latent dirichlet allocation. Journal of Machine Learning Research 3: 993–1022.

Carbonetto, P. and M. Stephens, 2012 Scalable variational inference for Bayesian variable selection in regression, and its accuracy in genetic association studies. Bayesian Analysis 7(1): 73–108.

Catchen, J., S. Bassham, T. Wilson, M. Cijrrey, C. O’Brien, Q. Yeates, and W. A. Cresko, 2013 The population structure and recent colonization history of Oregon threespine stickleback determined using restriction-site associated DNA-sequencing. Molecular Ecology 22: 2864–2883.

Engelhardt, B. E. and M. Stephens, 2010 Analysis of population structure: a unifying framework and novel methods based on sparse factor analysis. PLoS Genetics 6(9): e1001117.

Falush, D., M. Stephens, and J. K. Pritchard, 2003 Inference of population structure using multilocus genotype data: linked loci and correlated allele frequencies. Genetics 164(4): 1567–1587.

Hofman, J. M. and C. H. Wiggins, 2008 Bayesian approach to network modularity. Physical Review Letters 100(25): 258701.

Hubisz, M. J., D. Falush, M. Stephens, and J. K. Pritchard, 2009 Inferring weak population structure with the assistance of sample group information. Molecular Ecology Resources 9(5): 1322–1332.

Jordan, M. I., Z. Gharamani, T. S. Jaakkola, and L. K. Saul, 1998 An introduction to variational methods for graphical models. Machine Learning 37(2): 183–233.

Kadanoff, L. P., 2009 More is the same; phase transitions and mean field theories. Journal of Statistical Physics 137(5–6): 777–797.

Li, J. Z., D. M. Absher, H. Tang, A. M. Southwick, A. M. Casto, S. Ramachan-dran, H. M. Cann, G. S. Barsh, M. Feldman, L. L. Cavalli-Sforza, and R. M. Myers, 2008 Worldwide Human Relationships Inferred from Genome-Wide Patterns of Variation. Science 319(5866): 1100–1104.

Logsdon, B. A., G. E. Hoffman, and M. J. G, 2010 A variational Bayes algorithm for fast and accurate multiple locus genome-wide association analysis. BMC Bioinformatics 11(1): 58.

Mackay, D. J., 2003 Information theory, inference and learning algorithms. Cambridge University Press.

Novembre, J. and M. Stephens, 2008 Interpreting principal component analyses of spatial population genetic variation. Nature Genetics 40(5): 646–649.

Patterson, N., A. L. Price, and D. Reich, 2006 Population Structure and Eigenanalysis. PLoS Genetics 2(12): e190.

Pickrell, J. K. and J. K. Pritchard, 2012 Inference of population splits and mixtures from genomewide allele frequency data. PLoS Genetics 8(11): e1002967.

Price, A. L., N. J. Patterson, R. M. Plenge, M. E. Weinblatt, N. A. Shadick, and D. Reich, 2006 Principal components analysis corrects for stratification in genomewide association studies. Nature Genetics 38(8): 904–909.

Pritchard, J. K. and P. Donnelly, 2001 Casecontrol studies of association in structured or admixed populations. Theoretical Population Biology 60(3): 227–237.

Pritchard, J. K., M. Stephens, and P. Donnelly, 2000 Inference of population structure using multilocus genotype data. Genetics 155(2): 945–959.

Raydan, M. and B. F. Svaiter, 2002 Relaxed steepest descent and Cauchy-Barzilai-Borwein method. Computational Optimization and Applications 21(2): 155–167.

Reich, D., K. Thangaraj, N. Patterson, A. L. Price, and L. Singh. Reconstructing Indian population history. Nature 461(7263): 489–494.

Rosenberg, N. A., 2004 DISTRUCT: a program for the graphical display of population structure. Molecular Ecology Notes 4(1): 137–138.

Rosenberg, N. A., J. K. Pritchard, J. L. Weber, H. M. Cann, K. K. Kidd, L. A. Zhivotovsky, and M. W. Feldman, 2002 Genetic structure of human populations. Science 298(5602): 2381–2385.

Sato, M. A., 2001 Online model selection based on the variational Bayes. Neural Computation 13(7): 1649–1681.

Tang, H., J. Peng, P. Wang, and N. J. Risch, 2005 Estimation of individual admixture: analytical and study design considerations. Genetic epidemiology 28(4): 289–301.

Teh, Y. W., D. Newman, and M. Welling, 2007 A collapsed variational Bayesian inference algorithm for latent Dirichlet allocation. Advances in neural information processing systems 19: 1353.

Varadhan, R. and C. Roland, 2008 Simple and globally convergent methods for accelerating the convergence of any EM algorithm. Scandinavian Journal of Statistics 35(2): 335–353.

Wasser, S. K., C. Mailand, R. Booth, B. Mutayoba, E. Kisamo, B. Clark, and M. Stephens, 2007 Using DNA to track the origin of the largest ivory seizure since the 1989 trade ban. Proceedings of the National Academy of Sciences 104(10): 4228–4233.

